# Predicting Spatial Transcriptomics from H&E Image by Pretrained Contrastive Alignment Learning

**DOI:** 10.1101/2025.06.15.659438

**Authors:** Jiawei Zou, Kai Xiao, Zexi Chen, Jiazheng Pei, Jing Xu, Tao Chen, Likun Hou, Chunyan Wu, Yunlang She, Zhiyuan Yuan, Luonan Chen

**Author notes:** Corresponding authors: Luonan Chen, Zhiyuan Yuan, Yunlang She. These authors contributed equally to this work (Jiawei Zou, Kai Xiao).

## Abstract

The intricate molecular landscape within tissues holds crucial information about cellular behavior and disease progression, yet capturing this complexity at a spatial level remains challenging. While Spatial transcriptomics (ST) offers valuable insights into gene expression patterns within their native tissue context, its widespread adoption is hindered by high costs and limited gene detection capabilities. Here we introduce CarHE (Contrastive Alignment of gene expRession for hematoxylin and eosin image), a method that overcomes these limitations by accurately predicting high-dimensional ST data (over 10,000 genes) solely from readily available H&E (Hematoxylin and Eosin) stained images. This novel pre-trained architecture employs contrastive learning through two mechanisms: cell-type-based transcriptomics information transfer and image-based histology information transfer. These mechanisms precisely align image features with spatial single-cell gene expressions, achieving prediction accuracies exceeding 0.7 (up to 1.7 folds compared to second best) across diverse tissue types and species. CarHE’s superior performance extends to identifying subtle pathological features such as tertiary lymphoid structures in various cancers, including breast cancer, lung cancer, melanoma and ccRCC (clear cell renal cell carcinoma), and reconstructing 3D spatial transcriptomics from images alone, offering a cost-effective and robust alternative for large-scale spatial transcriptomics. We further validated CarHE’s effectiveness by predicting DFS (Disease-Free Survival) from >1,600 lung cancer patients HE images, achieving a significantly higher AUC (Area Under the Receiver Operating Characteristic Curve) of 0.73 compared to state-of-the-art alternatives (0.58-0.64).

## Introduction

Spatial transcriptomics (ST) has revolutionized our understanding of tissue biology by enabling the quantification of gene expression within its native spatial context, providing crucial insights into tissue development, disease progression, and three-dimensional organ architecture^1,2^. Despite its potential, current ST technologies, including imaging-based methods (e.g., Xenium^3,4^, MERFISH^5,6^, seqFISH^7-9^, STARmap^10^) and sequencing-based methods (e.g., 10x Visium^11^, Visium HD^11^, stereo-seq^12^), are constrained by cost, spatial resolution and/or gene coverage. Even advanced platforms like the Xenium, while offering single-cell resolution, remain cost-prohibitive and gene-limited.

The growing availability of large-scale spatial transcriptomics (ST) datasets—many with paired H&E images—has spurred computational methods to predict gene expression directly from histology. While methods like HisToGene^13^, Hist2ST^14^, ENLIGHT-DeepPT^15^, His2ST^14^ and BLEEP^16^ have emerged, they face fundamental challenges. The histology-to-expression relationship is inherently complex, and image features alone prove insufficient for comprehensive gene prediction. Critically, existing methods predict only a limited subset of genes (typically < 10,00). iSTAR^17^ and STEM^18^ leverages pre-trained H&E vision-transformer models to improve prediction accuracy. It further lack cross-tissue generalizability, that is, models trained on one tissue type exhibit significant performance degradation when applied to others.

These limitations are notable because pathologists readily identify major cell types from H&E images^19,20^, and cell types share common gene expression patterns across different tissues. Recent advancements in foundation models (e.g., scGPT^21^, scFoundation^22^, GMAI^23^ and Prov-GigaPath^24^) have provided powerful tools for feature extraction from both gene expression data and H&E images, providing an opportunity to address these limitations and develop a more comprehensive gene expression prediction solution.

Here, we present CarHE (Contrastive Alignment of gene expRession for H&E image), a pre-trained deep learning architecture that accurately and robustly predicts spatial transcriptomics profiles solely from H&E image across various tissue types. CarHE employs a two-step approach: first, a CLIP (Contrastive Language-Image Pre-Training)^25^ contrastive learning model aligns H&E image patches with cell type features, enabling reliable transfer of high-dimensional transcriptomics information through cell type associations. Second, an alignment refinement scheme using gradCAM^26,27^ maps image patches to single-cell features, facilitating precise spatial and histology information transfer. This approach enables comprehensive prediction of extensive gene expression profiles (over 17,000 genes) from H&E alone—surpassing gene coverage of current SOTA methods by >17-fold.

We first validated CarHE’s performance using the HER2ST (BRCA)^28^ and melanoma Xenium datasets, achieving prediction accuracies PCC (Pearson correlation coefficient) exceeding 0.7-0.8 and significantly outperforming existing methods by up to 2 folds across diverse tissue types and species. The model demonstrated utility in identifying tertiary lymphoid structures (TLS) in BRCA and ccRCC^29^ samples, detecting subtle features that pathologists might overlook. Furthermore, CarHE successfully reconstructed three-dimensional gene expression patterns from H&E images of the Dorsolateral Prefrontal Cortex (DLPFCdataset^30^, generating spatial structures that align well with established knowledge. To demonstrate cross-species applicability, we trained CarHE on mouse intestine 10X Visium HD data, achieving cell-type localization with an AUROC exceeding 0.9. The clinical relevance of CarHE was further validated using treatment outcome data from over 1,600 lung cancer patients. CarHE identified TLS precisely on these H&E slides and predicted DFS based on H&E images and imputed ST data with significant higher accuracy of 0.73 than exiting methods (0.58-0.64).

## Results

### The overview of CarHE

We constructed a comprehensive pretraining dataset comprising over 2 million cells or spots, each enriched with its spatial context, to facilitate downstream representation learning. To ensure broad generalizability across various sequencing technologies, the dataset integrates spatial transcriptomics data from four distinct assay platforms: Visium, Visium HD, ST (Spatial Transcriptomics), Xenium and Merfish (Figure 1A). This diverse corpus includes data spanning more than 10 human organs and tissues, originating from over 100 unique spatial slides derived from cancer samples, all accompanied by paired H&E-stained histology images. scGPT was applied to effectively learn a shared embedding space that harmonizes information across diverse sequencing protocols, bridging the gap between sequencing-based and imaging-based modalities. To facilitate downstream analysis, the embeddings of all cells and spots were clustered into 100 distinct clusters, which we refer to as “cell clusters.”

**Figure 1.**
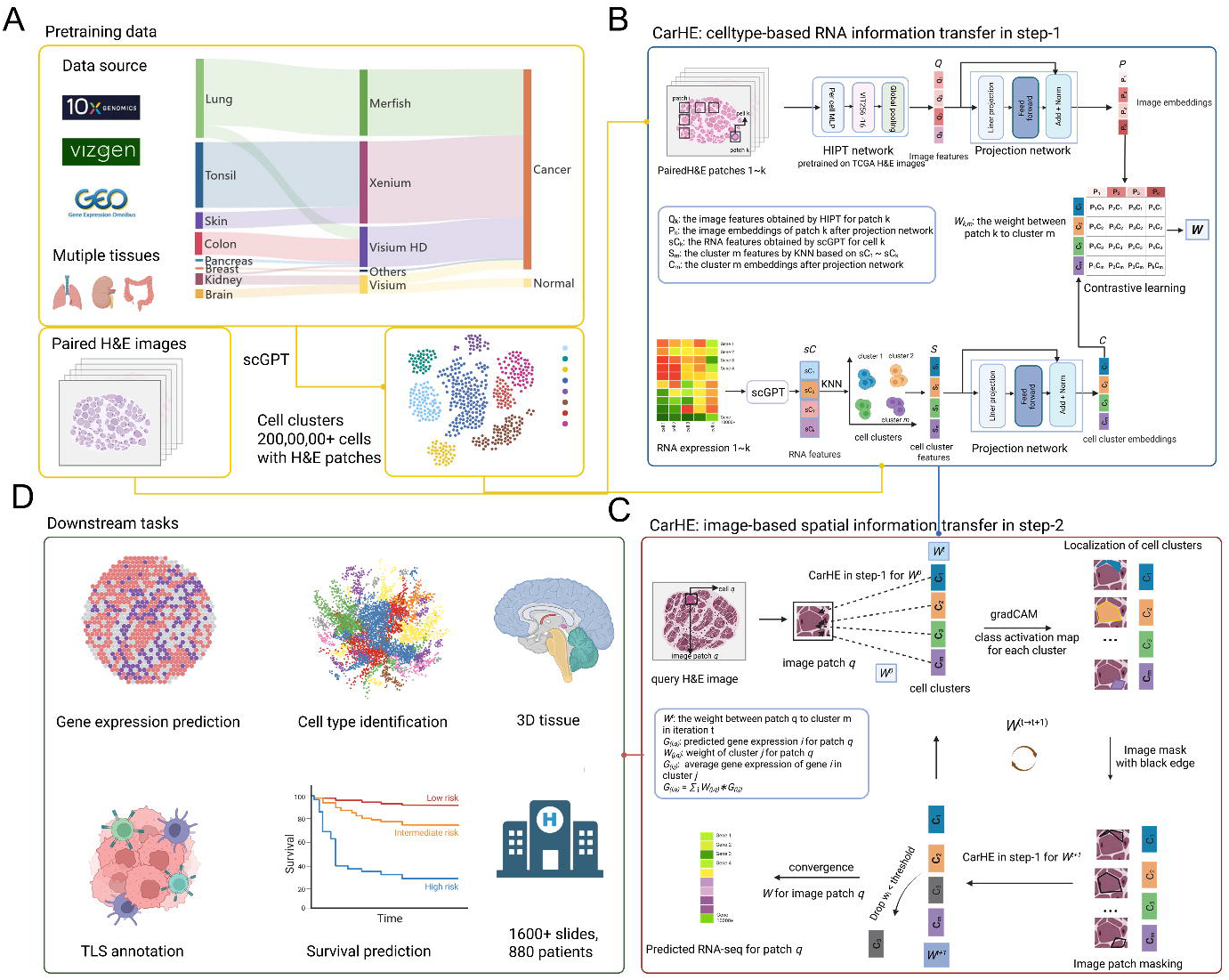
The overview of CarHE. A. The pretraining data of CarHE. B. The cell type based RNA information transfer of CarHE, leveraging HIPT and scGPT to extract features from H&E image and RNA-seq expression. The RNA-seq features are clustered as cell clusters and further aligned to paired H&E image after projection. The model results the weight *W* representing the match weight between image patches and cell clusters. C. The image-based spatial information transfer of to query H&E image patch *q*, the spatial location of each cell cluster represented RNA-seq is decided by an iterative refinement of weight *W*, based on marked image by grad-CAM. D. The downstream tasks of CarHE.

CarHE employs a novel two-step contrastive learning framework (Figure 1 B, C) that integrate both single-cell transcriptomics information and image-based spatial information to achieve comprehensive spatial transcriptomics prediction.

The H&E images were first segmented into image patches according to the location of spots or cells. That is, a patch *k, I*_*k*_ is an extended region centered around a spot or a cell with the size 256×256 pixels, about 0.5 μm/pixel. In the first step (cell-type-based RNA-seq information transfer), a Hierarchical Image Pyramid Transformer (HIPT) ^31,32^ model, pre-trained via self-supervised learning (SSL) on publicly available hematoxylin-and-eosin (H&E)-stained histology image datasets, is employed. HIPT extracts multi-scale image features (*Q*_k_) at two resolutions: 16x16 pixels (capturing fine-grained tissue characteristics) and 256×256 pixels (capturing broader structural features of the tissue) as one patch (taking patch-*k* as an example). Furthermore, the image features are reduced to image embedding features (*P*_k_) by a projection network. Simultaneously, gene expression features or RNA features (*sC*_k_) are extracted from the gene expression *x* for cell *k*, using a pre-trained single-cell foundation model (scGPT), specifically designed for RNA-seq data. For each spot or cell, the resulting embeddings are matched into one cell cluster S_m_, and the average feature of each cluster is used as its representation. Further, the cell cluster features are reduced to cell embedding features (C_m_). These gene expression embedding features (C_m_) of clusters are then aligned with H&E image embedding features (*P*_k_) via a modified contrastive learning approach inspired by CLIP^25^, which leverages a feed-forward neural network for projection (Figure 1A), i.e. obtaining the matrix *W*^*0*^ between cell clusters and image patches. *W*^*0*^ actually corresponds to the matrix between cell clusters and spots/cells, which is used as an initial matrix in the following second step, noting that each patch is centered on one spot. Training the model across diverse tissue datasets enables the transfer of RNA-seq-derived information from cell clusters to the corresponding cells in H&E images.

In the second step (image-based spatial information transfer), a circulating alignment model is developed to refine the spatial localization of cell clusters. For a given query image, cell clusters gene expression embedding features (C_m_) obtained in the pre-training data serve as a reference. Initial matrix or weights *W*^*0*^, representing the correspondence between cell clusters and the query image patch, are computed for the original H&E image. These weights *W*^*t*^ are iteratively (t **→** t+1) refined through recalculations based on edge-masked images generated using Grad-CAM^26,27^, progressively excluding unreliable matches. This iterative optimization results in the retention of only reliably matched cell clusters and their precise spatial locations, i.e. the converged matrix or weights *W* (Figure 1B). Thus, we obtain accurate reconstruction of spatial transcriptomics at a single spot or cell level from H&E-stained images alone by 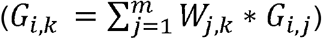.

Clearly, CarHE adopts two mechanisms: cell-type-based transcriptomics information transfer and image-based spatial information transfer. Through this dual-step approach, CarHE successfully bridges the gap between histological features and gene expression patterns, enabling accurate reconstruction of spatial transcriptomics data comprising over 10,000 genes from H&E-stained images alone.

Importantly, the pretraining strategy endows CarHE with robust gene expression prediction capabilities across diverse tissue types, while addressing the challenge of limited gene numbers. For any given H&E-stained histological image, CarHE enables accurate gene expression inference and precise cell type classification. Building upon these predictions, the model facilitates a range of advanced downstream applications, including 3D tissue reconstruction, identification of fine-grained biological structures, and survival outcome prediction. This comprehensive framework effectively integrates histological and genomic data, providing powerful tools for both exploratory and predictive tasks in spatial biology (Figure 1 D).

### Benchmarking of CarHE on BRCA ST Data and Melanoma Xenium Data

To benchmark CarHE’s accuracy in predicting spot-level gene expression, we validated the model performances on BRCA spatial transcriptomics (ST) data^28^, and melanoma Xenium data. The BRCA ST data from patient 1 and the melanoma Xenium data were removed from the training datasets. We predicted the gene expression from the H&E image of these two datasets using pre-trained CarHE, HisToGene, HisToST, THItoGene^33^ and STEM^18^, and compared the results with the ground truth. Since some of the methods were unable to predict large number of genes, we showed the results of top 3 genes i.e. CD3D, CD74 and COL3A1 (Figure 2 A). The prediction results of CarHE were consistent with manual annotations, demonstrating strong concordance between CarHE predictions and ground truth spatial distributions. To quantitatively assess performance, we calculated the Pearson correlation coefficient (PCC) between predicted and ground truth gene expression. The PCC results of CarHE prediction for the 3 genes were 0.64, 0.68 and 0.57, which were 1.5 folds higher than other methods. The overall PCC of top variable genes (300, 500, 1,000, 2,000) of CarHE prediction were 0.68, 0.70, 0.68 and 0.60, which were also significant higher than other methods (p value < 0.0001) (Figure 2. B). We also applied CarHE on an melanoma Xenium dataset. The prediction of cell type labels of CarHE were highly consistent with the ground truth, and the ARI (Adjusted Random Index) was 0.77. The detailed cell type prediction near the tumor border showed that CarHE could clearly identify the the CAFs (cancer associated fibroblasts) near tumor cells and T cells. Similar results were observed for ccRCC data prediction by CarHE (PCC = 0.65 for >17,000 genes; Supplementary Figure 1). The detailed benchmark results can be viewed in Table 1.

**Figure 2.**
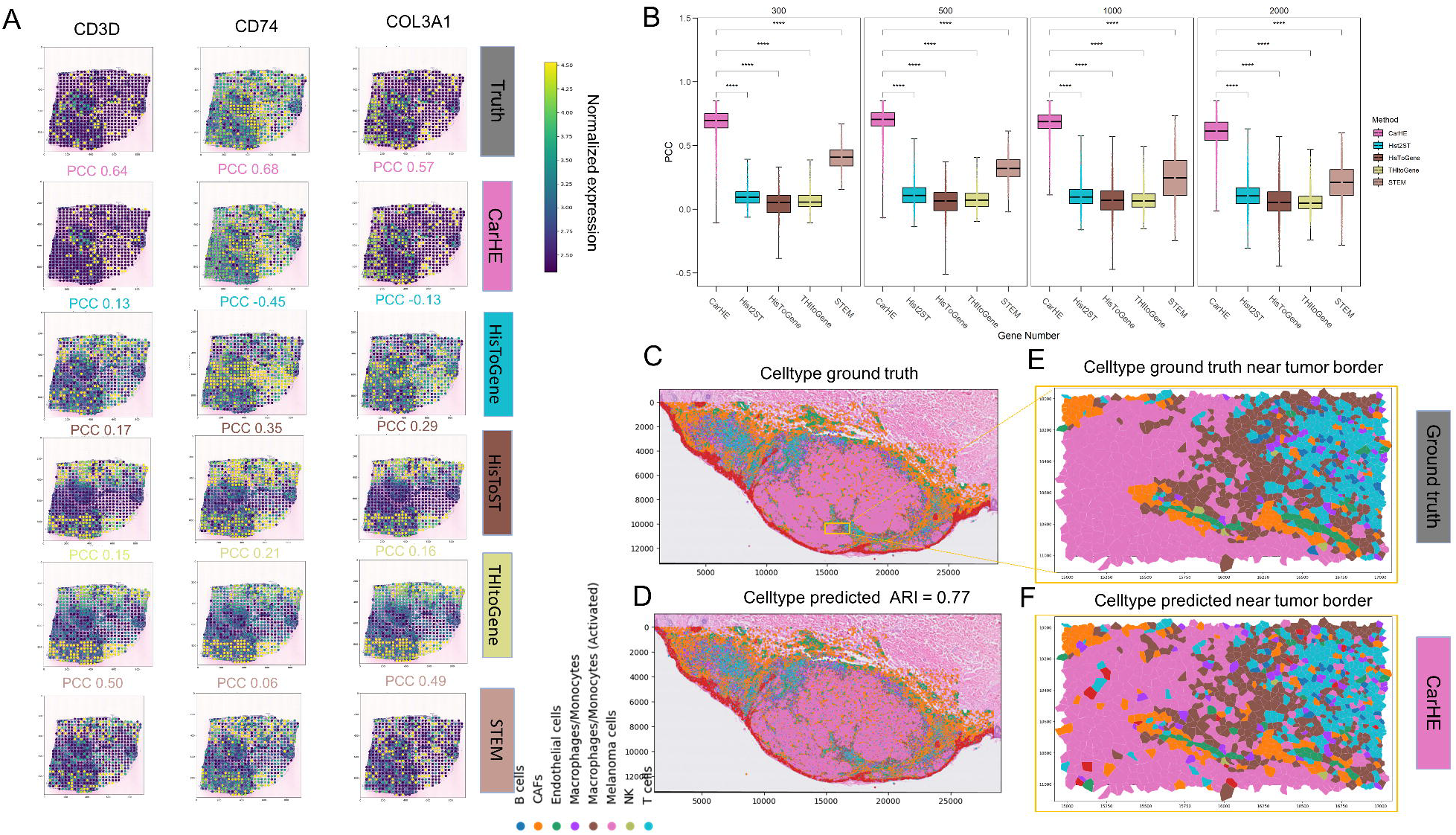
Benchmarking of CarHE on BRCA ST Data and Melanoma Xenium Data. A. Visualization of gene expression levels for CD3D, CD74, and COL3A1 overlaid on the H&E-stained image from a BRCA sample (patient 1). Results are illustrated for the ground truth and the predictions from four methods: CarHE, HisToGene, HisToST, THItoGene and STEM. B. Overall Pearson Correlation Coefficient (PCC) between predicted gene expression and ground truth across different methods. Results are separately shown for the top 300, 500, 1,000, and 2,000 most variable genes, highlighting CarHE’s performance relative to other approaches. C. Cell type annotations for the melanoma Xenium dataset, including a magnified view of detailed cell type distributions near the tumor border (highlighted within a yellow rectangle). D. Predicted cell type annotations for the melanoma Xenium dataset generated by CarHE, showcasing both the global cell type landscape and detailed predictions near the tumor border (yellow rectangle).

### CarHE identifies robust functional TLS area on cancer H&E images

To evaluate CarHE’s performance in identifying tertiary lymphoid structures (TLSs), we leveraged annotations from the original publication’s BRCA dataset^29,34^. Pathologists annotated TLSs (also known as ectopic lymphoid aggregates), important prognostic markers in cancer immunology, within the H&E images using color-coding (yellow for immune infiltrates; Figure 3A, E). Following the iSTAR method, TLS scores were calculated for patient F and H using the expression of known marker genes (i.e. CD3D, CD4, CD8A, CD79A; Supplementary Table 1). Figures 3B and 3C (patient 1) and Figures 3F and 3G (patient 2) show the TLS scores for both ground truth and CarHE predictions, exhibiting PCCs of 0.75 and 0.82, respectively. Comparison of high-scoring regions revealed strong concordance between CarHE predictions and manual annotations for patient 1 (Figure 3 B, C), except for one area initially annotated as cancer (Figure 3D). However, upon closer inspection (yellow dashed lines in Figure 3D), this area displayed immune infiltration features. In patient 2, CarHE predictions accurately identified both manually annotated immune infiltrated areas (Figure 3. F, G). Although two small cancerous regions showed high ground truth TLS scores in the second area (Figure 3H), CarHE’s lower scores in these regions better reflected the H&E image features characteristic of cancerous tissue. This suggests that high TLS gene expression in these ground truth spots might reflect signal bleed-through from neighboring immune cells. Overall, these results demonstrate that CarHE accurately predicts TLS features consistent with both ground truth annotations and H&E image characteristics.

**Figure 3.**
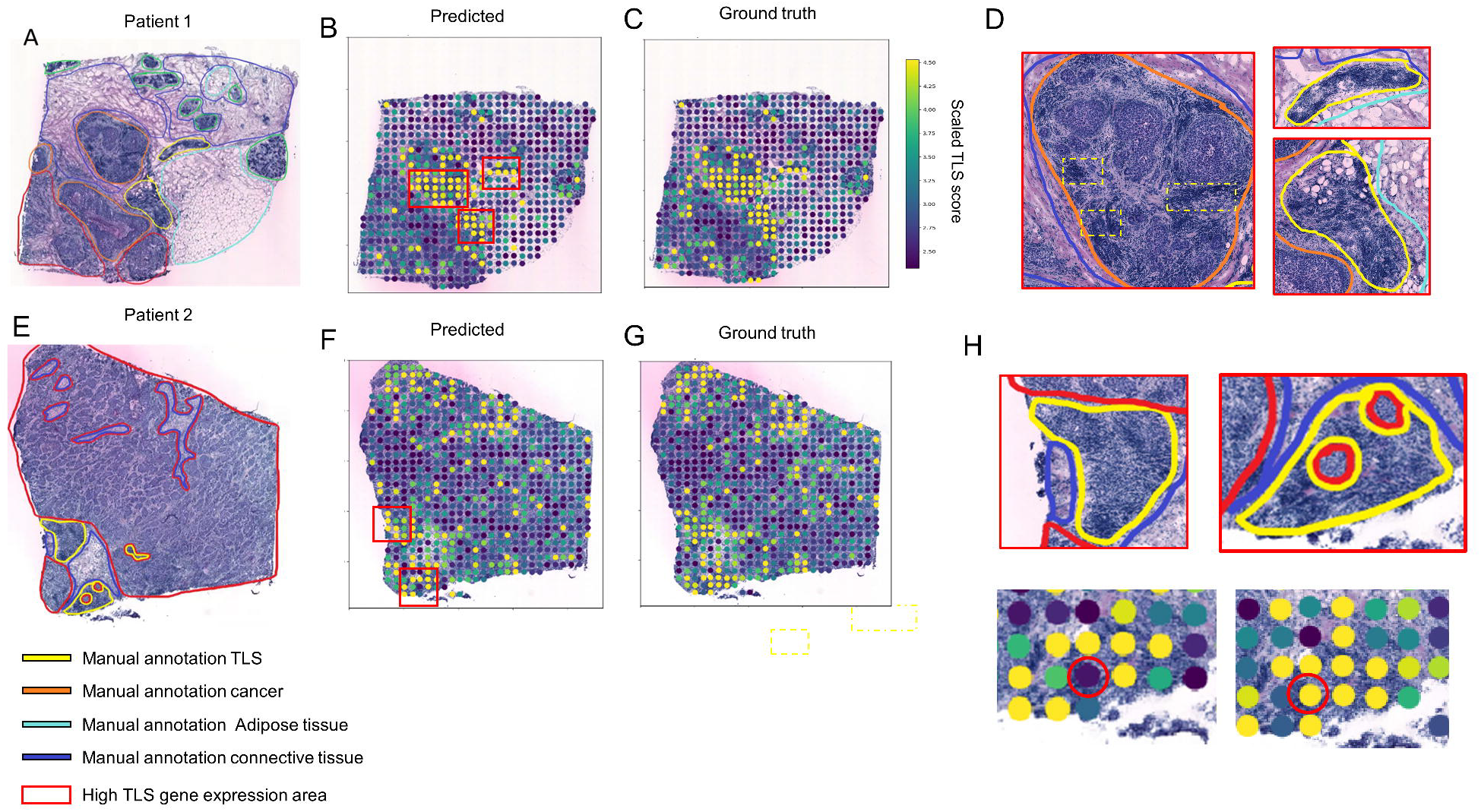
The TLS areas identified by CarHE prediction. A. The manual annotation of H&E slide of patient 1, whereas yellow refers to immune infiltrated area; red refers to cancer area; light-blue refers to adipose area; blue refers to connective area. B. The TLS score calculated by CarHE prediction of patient 1, whereas red rectangle represents three areas that might be TLS. C. The TLS score calculated based on the ground truth. D. The detailed manual annotation of the three TLS areas predicted by CarHE. E. The manual annotation of H&E slide of patient 2. F. The TLS score calculated by CarHE prediction of patient 2. G. The TLS score calculated based on the ground truth.H. Red rectangle represents the detailed manual annotation of the TLS area. Below are the detailed TLS score predicted by CarHE and the ground truth of, whereas red circle represents a cancer area.

### CarHE constructs 3D structure of DLPFC solely by H&E images

To evaluate CarHE’s capability in reconstructing complex 3D tissue architecture solely from H&E images, we analyzed the DLPFC dataset^30^, focusing on four consecutive tissue sections (151673, 151674, 151675, and 151676; Figure 4A-E). While the predicted gene expression patterns showed moderate correlation with ground truth measurements (Pearson correlation coefficient (PCC) =0.32; Supplementary Figure 2), this lower correlation comparison to cancer datasets likely reflects the HViT model’s initial pre-training bias toward TCGA cancer specimens rather than normal tissues.

**Figure 4.**
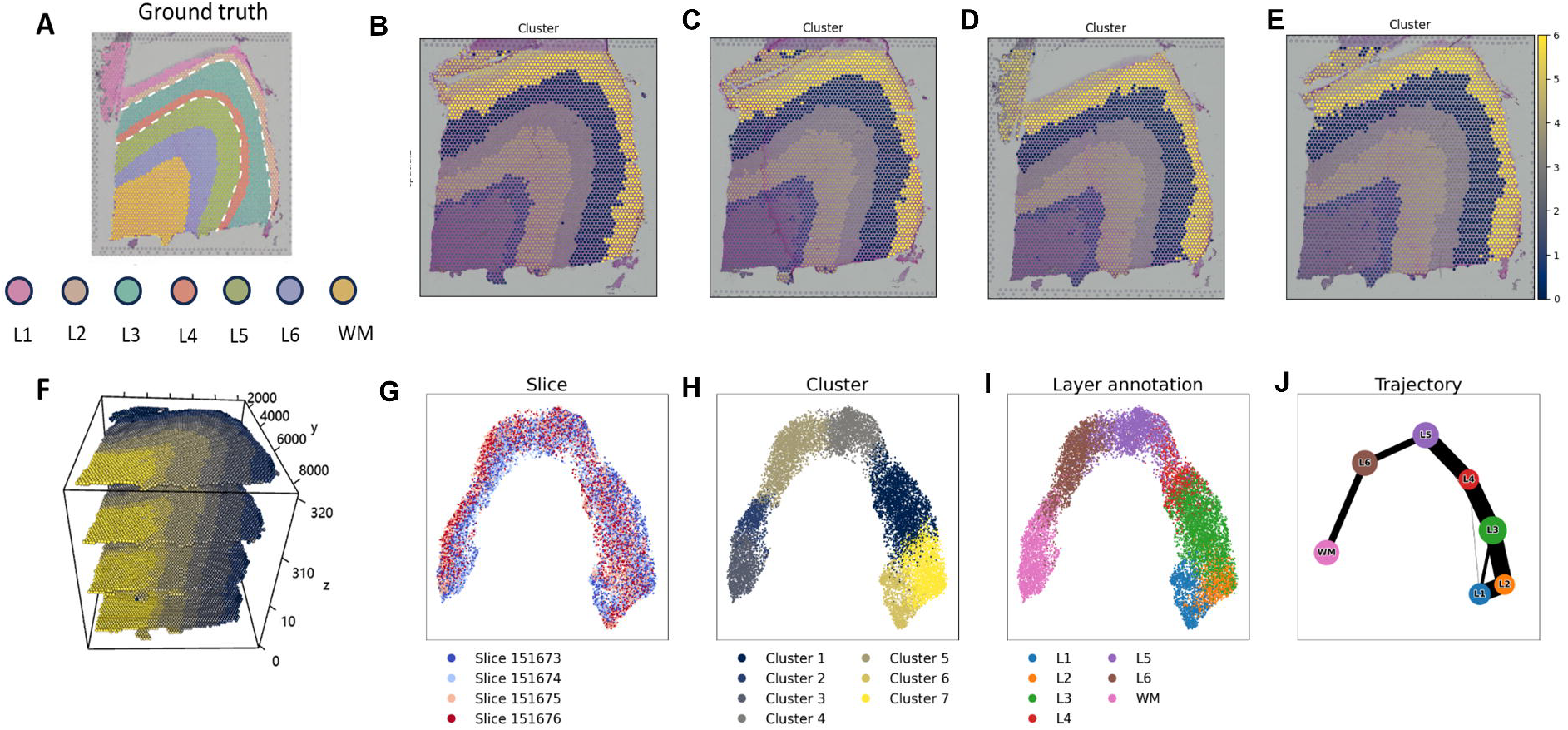
The application of CarHE on human DLPFC dataset. A. the ground truth of the structure layers of DLPFC ST dataset. B-E the k-means cluster based on CarHE prediction only by H&E images, from slide 151673 to 151676. F. the 3D structure of CarHE prediction. G. The slice distribution on umap by CarHE prediction. H. The umap cluster. I. The manual layer annotation on umap. J. The trajectory reference using PAGA, by CarHE prediction.

Despite this limitation, simple k-means clustering (k=7) of the predicted gene expression profiles revealed spatial patterns that closely matched expert manual annotations of cortical layers (Figure 4A-E). The three-dimensional reconstruction using Stitch3D further validated these findings, demonstrating clear preservation of the laminar organization characteristic of cortical architecture (Figure 4F).

To understand the developmental implications of this spatial organization, we performed trajectory analysis using UMAP dimensionality reduction coupled with PAGA (Figure 4G-J). The resulting trajectory map revealed a clear progression from layer 1 through layer 6 and into the white matter (WM), accurately reflecting the established developmental sequence of cortical layer formation.

### CarHE predicts precise cell type distribution on mouse dataset

CarHE was also applied to a mouse intestinal 10x Visium HD dataset, where gene expression was measured in 2x2 µm patches. Initially, StarDist^35,36^ was used for cell segmentation, generating a single-cell resolution spatial transcriptomics (ST) dataset. CarHE was trained on the right half of the dataset (coordinates > 50,00 µm), with the left half used for testing. Due to the dataset-specific training, the number of cell clusters was reduced to 21, corresponding to the number of clusters identified using the Leiden algorithm (resolution = 1). Predicted and ground truth cluster classifications are shown in Figure 5 A, B. Focusing on marker genes for enterocytes, goblet cells, Paneth cells, and T cells, we compared the predicted and ground truth spatial distributions of these cell types (Figure 5C). The area under the receiver operating characteristic curve (AUROC) exceeded 0.9 for all cell clusters (clusters). Given the functional divergence between intestinal villi and glands, we further analyzed the spatial distribution of *Lyz1*, a marker gene primarily expressed in the glands. As shown in Figure 5D, E, both predicted and ground truth *Lyz1* expression were predominantly localized to the gland region, with minimal expression in the villi, highlighting CarHE’s robust performance across diverse tissue types and data modalities.

**Figure 5.**
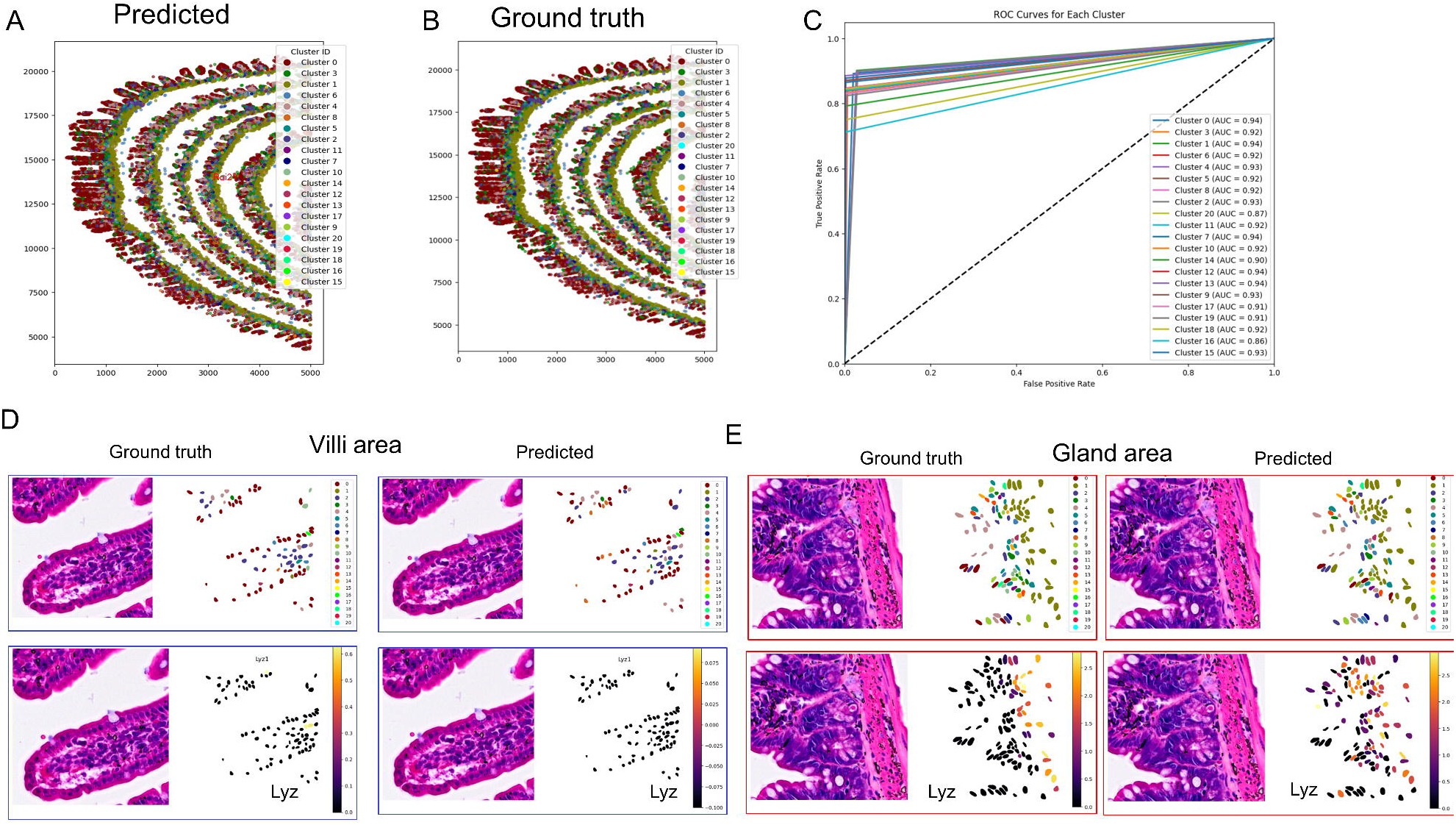
The prediction of the mouse intestine visium HD dataset. A. The distribution of predicted cell clusters by CarHE. B. The distribution of cell clusters of the ground truth. C. The distribution of four main cell types. D. The AUROC of prediction of cell clusters. D. The cell cluster distribution and gene expression of Roi 1 of ground truth and prediction locate in the villi area. E. The cell cluster distribution and gene expression of Roi 1 of ground truth and prediction locate in the gland area.

### CarHE identifies TLS precisely on lung H&E slides and predict accurate DFS for patients

As an application of CarHE, we evaluated its performance on over 1,600 hematoxylin and eosin (H&E)-stained lung cancer slides obtained from 881 patients (in-house dataset). Spatial transcriptomics (ST) data were imputed at the spot-level across these slides, capturing the cellular composition and TLS (tertiary lymphoid structure) scores. Among the cohort, cancer regions and TLS areas in 62 slides were manually annotated by pathologists. To assess the accuracy of CarHE, we first compared regions with high TLS scores (spots exhibiting statistically significant TLS scores above the median of other spots) predicted by CarHE with manual annotations (Figure 6A, B). The results demonstrated strong concordance, as the predicted TLS regions (marked by blue rectangles) closely aligned with the manually annotated TLS areas (highlighted by green circles).

**Figure 6.**
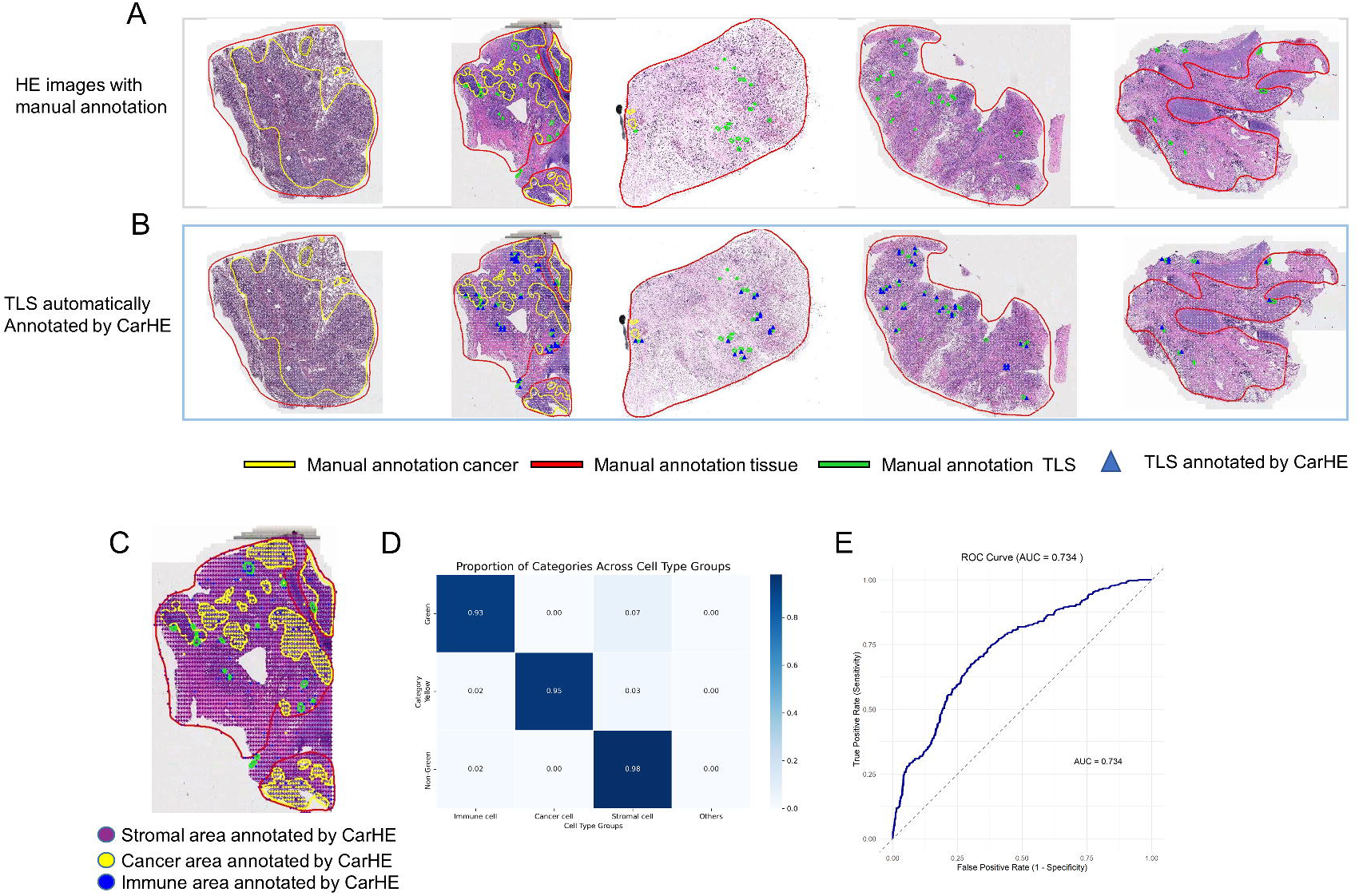
Application of CarHE on in-house lung cancer datasets. A Manual annotation of H&E slides by pathologists. Red circles delineate overall tissue contours; yellow circles represent cancer areas; green circles denote TLS (tertiary lymphoid structure) areas. B Automated TLS annotation performed by CarHE. Blue rectangles indicate TLS regions identified by CarHE, showcasing alignment with manual annotations. C Distribution of cell types predicted by CarHE across annotated areas. Purple spots represent stromal cells as identified by CarHE; yellow spots indicate cancer cells; blue spots correspond to immune cells, including B cells, T cells, and macrophages. D Heatmap illustrating the accuracy of CarHE cell type predictions within the three manually annotated categories (cancer areas, stromal regions, and TLS regions). High concordance levels were observed between CarHE predictions and manual annotations. E CarHE’s performance in predicting disease-free survival (DFS) for the validation patient cohort. The model integrates features derived from image patches and predicted gene expression, yielding robust predictive performance.

Additionally, we analyzed the primary cell type distributions in these regions and presented the findings in Figure 6C. Consistent with expectations, the cancer areas were predominantly identified as cancer cells, whereas the TLS regions were primarily imputed as immune cell populations, including B cells, T cells, and macrophages. The correspondence between cell types inferred by CarHE and the three manually annotated categories demonstrated high concordance, with correct annotations exceeding 93%, 95%, and 98% for the respective categories (Figure 6D).

Finally, we trained a simple multi-layer perceptron (MLP) model to predict disease-free survival (DFS) for patients. This model utilized both image patch features and the inferred gene expression features generated by CarHE. On the validation cohort, the MLP model achieved an area under the curve (AUC) of 0.734, substantially outperforming state-of-the-art methods, which reported AUCs of 0.586 and 0.643, respectively. This represents a 15% improvement in predictive performance relative to existing approaches.

## Discussion

In this study, we introduced CarHE, a pre-trained deep learning framework designed to predict high-dimensional spatial transcriptomics (ST) data solely from H&E-stained images. The method leverages a two-step contrastive learning approach to align histology image features with cell-type-resolved gene expressions. The first step establishes connections between image patches and cell type features using a CLIP-based contrastive learning model, while the second step refines these mappings with gradCAM to ensure precise spatial and histological information alignment. This innovative framework effectively addresses key limitations in spatial transcriptomics, enabling predictions of gene expression patterns across diverse tissue types and species with exceptionally high accuracy.

CarHE offers several distinct advantages. First, it significantly reduces costs and expands accessibility by eliminating the need for expensive and complex ST technologies. Second, it achieves high predictive accuracy (>0.83 Pearson correlation coefficient), outperforming existing methods by up to two-fold and extending predictions to over 17,000 genes. Third, the model demonstrates impressive generalizability across tissues and species, as evidenced by its performance with datasets from human cancers, mouse intestine, and the human dorsolateral prefrontal cortex dataset (DLPFC). Moreover, it provides critical insights into subtle pathological features, such as tertiary lymphoid structures (TLS), and reconstructs complex three-dimensional transcriptomic profiles from images alone. In a practical application, CarHE outperformed other approaches in predicting disease-free survival (DFS) from >1,600 lung cancer patient H&E images, achieving a significantly higher AUC of 0.73 compared to traditional methods.

Despite the promising outcomes, several limitations remain and warrant further investigation. First, the predictive performance relies on the quality of input H&E images and ST data, with robustness to low-quality datasets still needing improvement. Second, while the method generalizes across various tissue types, its performance across rare or underrepresented tissue types requires further validation. By training on larger amount of data, the robustness of CarHE could be improved in future.

In conclusion, CarHE represents a groundbreaking advancement in integrating spatial transcriptomics and histological image-based gene expression prediction. It addresses significant cost and scalability challenges while providing accurate and robust predictions across diverse biological and clinical scenarios. Continued research should focus on improving robustness, refining cross-tissue generalizability, and optimizing computational efficiency to unlock its full potential for widespread adoption.

## Methods

### Preprocessing of scRNA and images

#### H&E image preprocess

Before training, the H&E images were first normalized with staintools, using “LuminosityStandardizer” to eliminate the difference between the stains of different tissue slides. Then the H&E images were segmented into image patches according to the location of spots or cells. The size of each patch is 256×256 pixels, with about 0.5 µm/pixel.

#### Gene expression preprocess

To eliminate the batch effect between gene expression of different datasets and technologies, we first merged all datasets together with scanpy. After quality control, whereas spots or cells with less than 100 genes or transcripts expressed were filtered. Then, the top variable genes were detected by “scanpy.pp.highly_variable_genes” and used later in scGPT.

#### scGPT

The single-cell foundation model scGPT is used for RNA feature extraction, whereas the pre-trained model on 33 million human cells is applied. The input for the scGPT model is the matrix gene expression of each cell which contains up to 10,000 highly variable genes detected by “scanpy.pp.highly_variable_genes”. The output of the cell embedings is the 512-dimention RNA features for each cell.

#### HIPT

We apply HIPT model to extract image features for each 256×256 image patches. The HIPT model first applied a VIT model (n=8,h=6,d=384) to extract feature at 16 pixel level. The output is the 384-dimention features of each 16 pixel mini-patch within the 256×256 image patch. Then these features were input into another VIT model (n=4,h=6,d=192) to get the 256×256 patch level features. The HIPT model was pre-trained on TCGA H&E slides. Here, we fine-tuned it with our training datasets derived various cancer slides, by training the last two transformer block while keep the others frozen.

#### Projection network

The projection network is a neural network module. It receives an input embedding of dimension (384 for image, 512 for gene expression). The network then projects this input to a projected feature with 256 dimensions using a linear layer. This is followed by a GELU non-linearity and another linear layer that maintains the 256 dimensions. Dropout is applied for regularization. A key feature is the residual connection, which adds the output of the initial linear projection (the 256 dimensions output) to the result after the second linear layer and dropout. Finally, layer normalization is applied, again operating on the 256 dimensions features. The module’s output is a tensor of dimension 256. This architecture, effectively an MLP with a residual connection and layer normalization, aims to create a representation space where distances between these 256-sized embeddings are more meaningful for the contrastive loss function, thus facilitating effective unsupervised feature learning.

### CarHE (Contrastive Alignment of gene expRession for hematoxylin and eosin image)

#### Step-1: cell-type-based RNA-seq information transfer

In the first step of CarHE (cell-type-based RNA-seq information transfer), contrastive learning is employed to model the correspondence between histological H&E image features and cell-type-specific gene expression features.

After preprocess of datasets, the model is trained with paired H&E images and corresponding gene expression data, yielding feature vectors for each modality. For H&E image patch *I*_*k*_, *I*_*k*_ ∈ ℕ ^3×256×256^, we also have the paired gene expression *x* for cell *k, x*_*k*_ ∈ ℕ^*gene*^.

For H&E patch *I*_*k*_, the HIPT model is utilized to extract features at both local (16x16 pixels) and global (256×256 pixels) scales, leveraging self-supervised learning pre-trained on The Cancer Genome Atlas (TCGA) dataset.

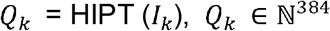

Later, the projection network projected the *Q*_*k*_ to a lower dimension of 256.

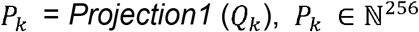

For gene expression profiling *x*_*k*_, the scGPT foundation model, pre-trained on large-scale single-cell RNA-seq datasets, is applied to encode gene expression embeddings.

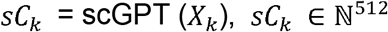

These embeddings *sC*_*k*_, derived from scGPT across samples, are grouped into a predefined number (m) of cell clusters (default = 100) using K-nearest-neighbors clustering. Each spot’s or cell’s individual gene expression embedding is subsequently replaced with the mean embedding of the cell cluster to which it belongs, providing a simplified and biologically relevant representation.

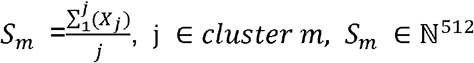

Later, another projection network projected the *S*_*m*_ to a lower dimension of 256.

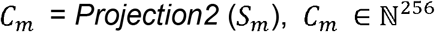

A pairwise weight is then established between H&E image features and semi-cell-type gene expression embeddings, aligning them through a modified contrastive learning paradigm. This alignment facilitates the transfer of transcriptomic information to histological contexts, representing the first step of cross-modality integration in the framework. The model is trained with paired H&E images and corresponding gene expression data, yielding feature vectors for each modality. The contrastive loss is:

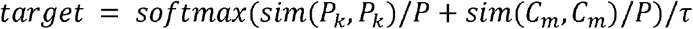

where,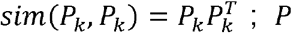; is a symmetric matrix with diagonal elements of 1 and a constant value *α* elsewhere,,*α* > 1;*τ* is a learnable temperature hyperparameter.

The final contrastive loss is:

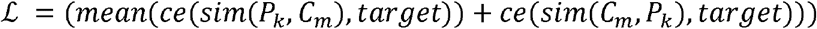

where,*ce* (*cross ropy*) is a cross-entropy loss.

By these procedures, we can get a weight *w*_*k*,*m*_ between any image patch *I*_*k*_ and cell cluster *C*_*m*_.

#### Step-2: image-based spatial information transfer

The second step employs an iterative scheme for gene expression prediction and spatial mapping of those expressions within the H&E image. Given a target H&E image, denoted by patch *q*, and a set of candidate cell clusters *C*_*m*_, an iterative process is initiated. First, initial matching weights *w*^0^ between the target H&E image *q* and each cell clusters *C*_*m*_ are computed using the trained CarHE in step 1. Each gene expression is treated as a class, and Class Activation Maps (CAMs) are generated using Grad-CAM to identify potential spatial regions. These initial CAM proposals are often noisy; they can be enhanced visually prompted images, through the application of visual cues of black edges to highlight identified regions. These visually enhanced images are then used to compute refined matching weights *w*^*t*^. Any gene expression category with a matching probability below a threshold of 0.4 is discarded. This refinement process is iteratively repeated, with subsequent visual cues and matching probability recalculations, until the predicted set of gene expressions and their spatial distributions within the image reach convergence (*w*^t^ = *w*^*t*-1^). The converged output *w*^*t*^, reflects the final gene expression prediction and spatial distribution within the H&E image. And the predicted gene expression *G*_*q*_ for patch *q* can be write as:

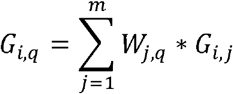

Where *w* _*j*,*q*_ is the weight of cluster *j* for patch *q, G*_*i*,*j*_, is the average gene expression of gene *i* in cluster *j*.

##### Algorithm image-based spatial information transfer.

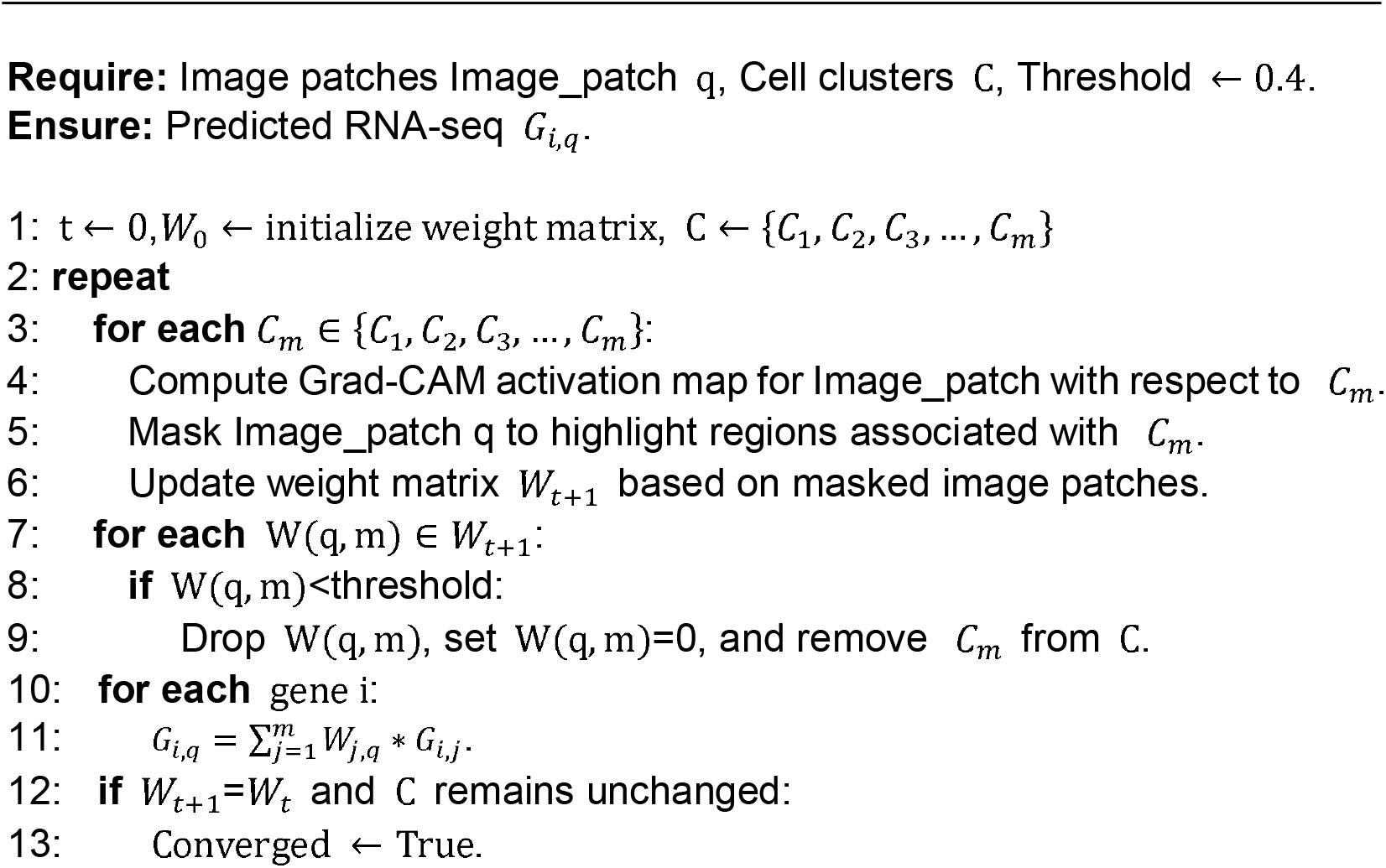
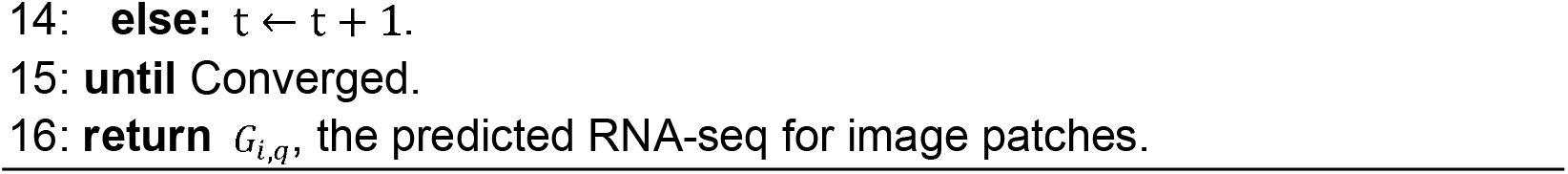

### Baseline methods

HisToGene^13^ adopted a ViT as the image encoder, leveraging the self-attention mechanism to extract global features, which were subsequently projected onto the gene expression space through fully connected layers.

His2ST^14^ integrated a Convmixer module to capture local 2D visual patterns, a Transformer module to model global spatial dependencies via self-attention, and a GNN module to explicitly learn neighborhood relationships between spots.

THItoGene^33^ used dynamic convolutional and capsule networks to extract molecular-relevant signals from H&E-stained histological images.

Stem^18^ applied a conditional diffusion model that utilizes DiT blocks and pathology foundation models to iteratively generate gene expression profiles conditioned on image patches from H&E-stained slides.

### Datasets

The pre-training data includes spatial transcriptomics data from four distinct assay platforms: Visium, Visium HD, ST (Spatial Transcriptomics), and Xenium. The Visium datasets includes one DLPFC dataset^30^ and one ccRCC dataset^29^. The DLPFC dataset (https://github.com/LieberInstitute/spatialDLPFC) contains 12 slides, and slides 151673, 151674, 151675, 151676 are kept for testing. The ccRCC dataset contains 23 slides, whereas slides c2, c3 and c4 are kept for testing. The Visium HD dataset were derived from multiple cancers, including lung cancer, colorectal cancer, kidney cancer and pancreas cancer. These datasets are collected from 10 x genomics website (https://www.10xgenomics.com/datasets). The ST datasets refers to the BRCA dataset (https://github.com/almaan/her2st), which contains 36 slides, the test slide is slide H1. The Xenium datasets contains various cancer slides, i.e. lung cancer, breast cancer, colon cancer and tonsil dataset. The melanoma dataset is kept for testing. These datasets are collected from 10 x genomics (https://www.10xgenomics.com/datasets). The WSI dataset of 880 lung cancer patients was sourced from four medical centers in China: Shanghai Pulmonary Hospital, The First Affiliated Hospital of Nanchang University, Ningbo HuaMei Hospital, and The First Hospital of Lanzhou University.

## Code availability

The CarHE algorithm was implemented in Python and is available on GitHub at https://github.com/Jwzouchenlab/CarHE.

## Acknowledgements

We thank all the members of Chen laboratory for technical assistance. This work has been supported by National Key R&D Program of China (2022YFA1004800, 2025YFF1207900), Natural Science Foundation of China (T2341007, T2350003, 12131020, 42450084, 42450135, 12326614, and 12426310), Science and Technology Commission of Shanghai Municipality (23JS1401300), Zhejiang Province Vanguard Goose-Leading Initiative (2025C01114), Hangzhou Institute for advanced study of UCAS (2024HIAS-P004), and JST Moonshot R&D (JPMJMS2021).

## Authors’ contribution

Jiawei Zou developed the algorithm and drafted the manuscript. Kai Xiao performed benchmarking of the methods and developed the code. Zexi Chen, Jiazheng Pei, and Jing Xu contributed to discussions and provided insights that enhanced the development of the method. Tao Chen processed the in-house data and contributed to the improvement of the manuscript. Likun Hou and Chunyan Wu annotated the in-house slides. Yunlang She managed the in-house data and contributed to manuscript writing. Zhiyuan Yuan refined the method, enhanced the figures, and co-authored the manuscript. Luonan Chen supervised the project, designed the algorithm, and provided substantial contributions to manuscript writing.

## Legends

Supplementary Figure 1. The prediction of ST by CarHE on ccRCC H&E images.

Supplementary Figure 2. The prediction of ST on human DLPFC slides. A-D. The ground truth expression of top 4 variable genes of ground truth. E-H. The gene expression of CarHE prediction.

Table 1. Benchmark of CarHE and other methods on ST data.

Supplementary Table 1. The genes used to identify TLS.

