## Supplementary figures and images for "Predicting Spatial Transcriptomics from H&E Image by Pretrained Contrastive Alignment Learning"

### Supplementary Figure 1

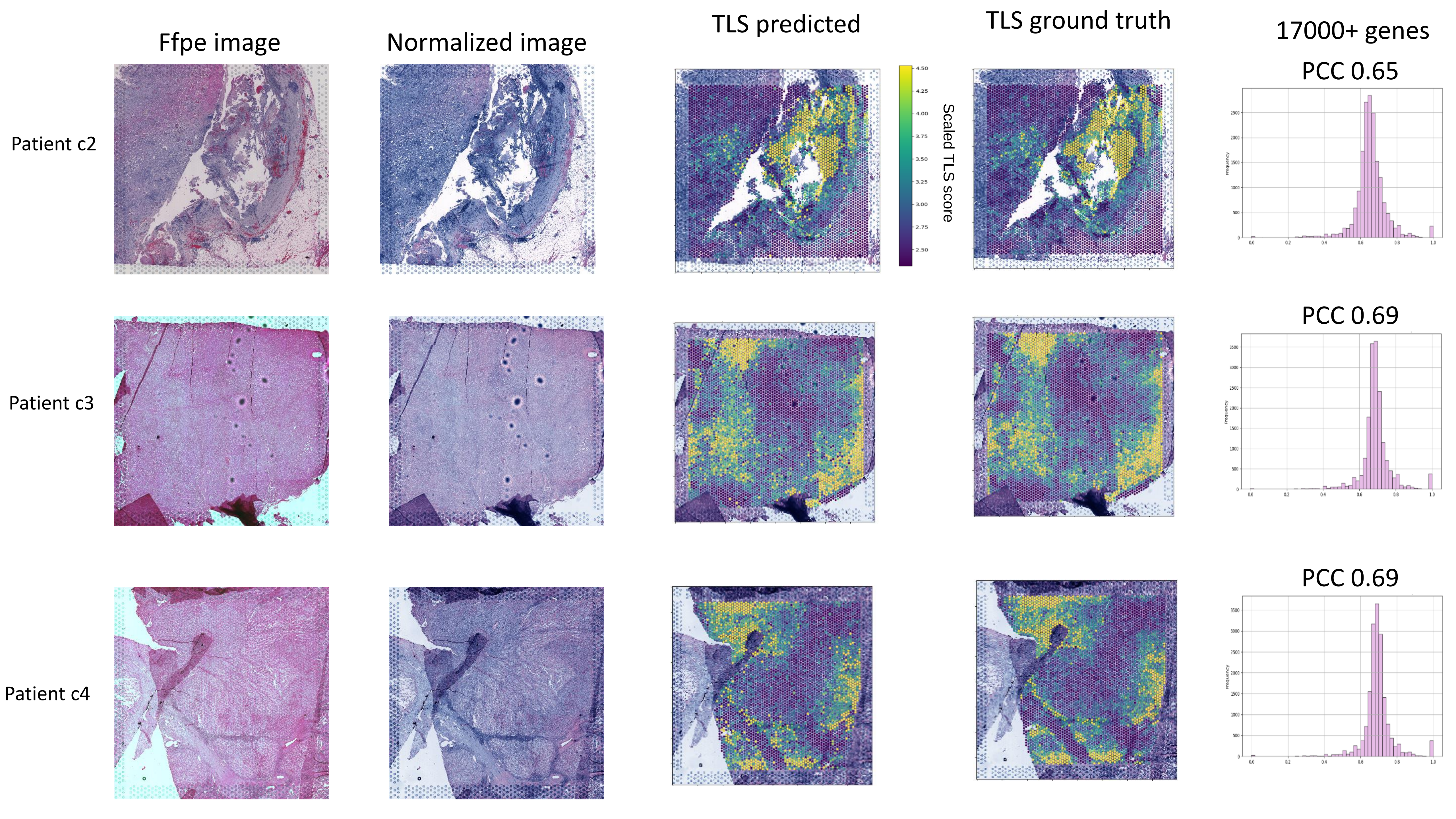

### Supplementary Figure 2

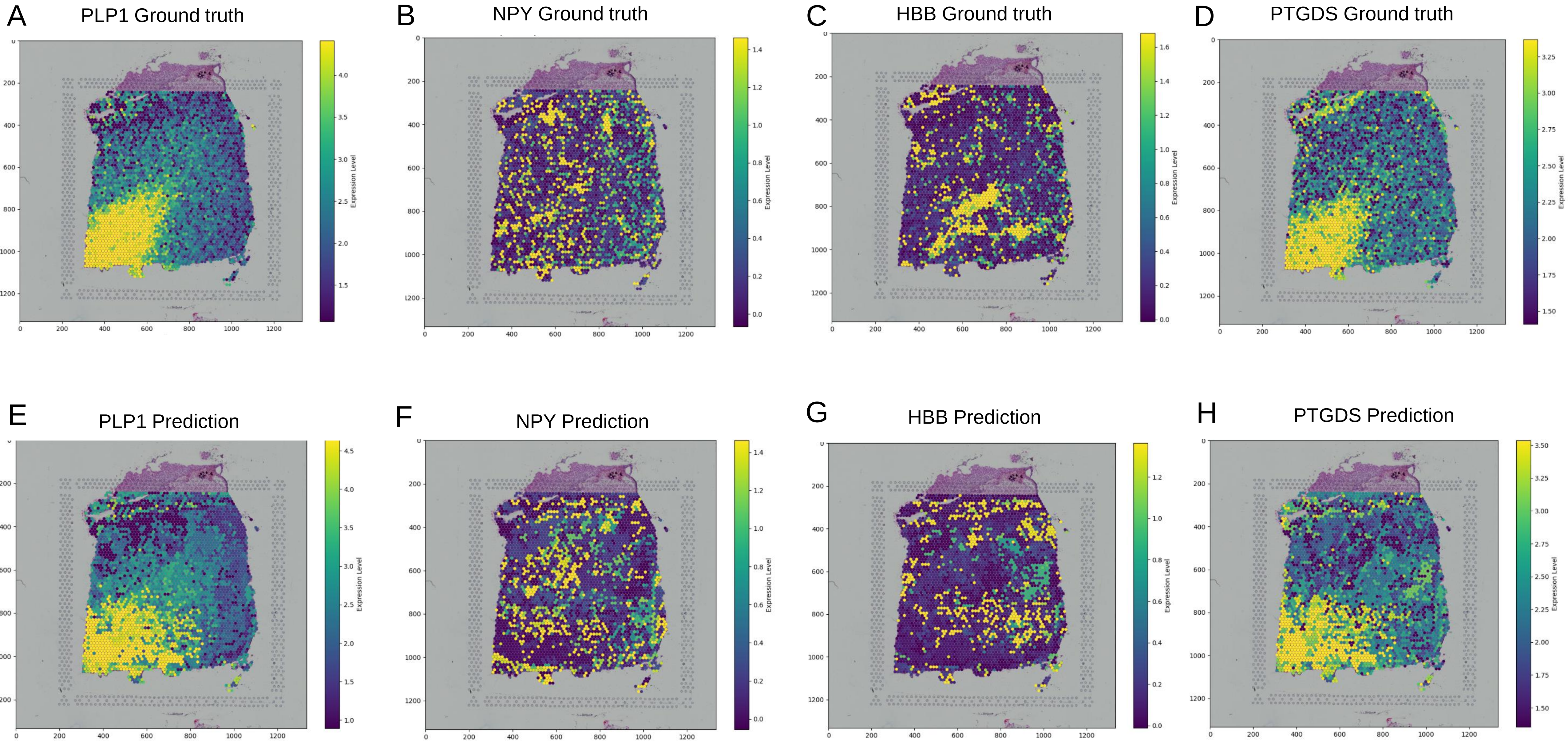
